# GPU accelerated population genetics statistics using pg_gpu

**DOI:** 10.64898/2026.05.29.728868

**Authors:** Nathaniel S. Pope, Angel G. Rivera-Colón, Ananya Kapoor, Kevin Korfmann, Murillo F. Rodrigues, Scott T. Small, Anastasia A. Teterina, Andrew D. Kern

## Abstract

Population genetics summary statistics—diversity, divergence, linkage disequilibrium, selection scans, and dimensionality reduction—are fundamental across human, agricultural, and ecological genomics. As whole-genome sequencing datasets have grown to hundreds of thousands of individuals, the cost of computing these statistics on conventional CPU implementations has become a major bottleneck: windowed scans of a single chromosome arm can take hours to days, and computation of pairwise linkage-disequilibrium statistics useful for demographic inference scales as *O*(*n*^2^) in sample size, often exceeding wall-clock budgets entirely. We present `pg_gpu`, a Python library implementing a comprehensive catalog of population-genetics summary statistics as fused CUDA kernels on NVIDIA GPUs. `pg_gpu` covers eleven categories spanning diversity and neutrality tests, divergence, admixture, the site-frequency spectrum, linkage disequilibrium, haplotype-based selection scans, dimensionality reduction (PCA, randomized PCA, local PCA / `lostruct`), distance distributions, relatedness, resampling, and a generalized weighted-SFS framework for custom *ω* estimators. On the full Ag1000G Phase 3 chromosome 3R arm (2,940 haplotypes, 10.9 million variants) `pg_gpu` agrees with `scikit-allel` and `PLINK2` to machine precision while delivering a median 139× and maximum 1,096× speedup. For the multi-population LD statistics used by `moments` for demographic inference, `pg_gpu` is a drop-in replacement that yields a ~1,750-fold speedup over the native implementation. Whole chromosome arm scans, `lostruct` screens, and calculation of LD statistics complete on a single NVIDIA A100 in seconds to a few minutes.

## Introduction

Population genetics relies on a rich toolkit of summary statistics to characterize genetic variation within and between populations. Statistics such as nucleotide diversity (*π*), Tajima’s *D, F*_*ST*_, and linkage disequilibrium (LD) are foundational to studies of demographic history, natural selection, population structure, and admixture. As whole-genome sequencing datasets have grown from hundreds to hundreds of thousands of individuals—exemplified by projects such as the Anopheles 1000 Genomes (Anopheles gambiae 1000 Genomes Consortium 2017) and UK Biobank (Bycroft et al. 2018)—the computational cost of these analyses has become a serious bottleneck.

A number of software tools have been developed to compute population genetics statistics from genomic data. `PLINK` (Purcell et al. 2007; Chang et al. 2015) was the first high-performance tool to offer a broad range of genome-wide association as well as other simple population-based summaries. `VCFtools` (Danecek et al. 2011) was a general-purpose tool that provided command-line access to diversity, *F*_*ST*_, and LD calculations from `VCF` files, but was designed for the sample sizes of the early 1000 Genomes era. `ANGSD` (Korneliussen, Albrechtsen, and Nielsen 2014) is a high-performance C/C++ tool, but it is oriented towards genotype likelihood workflows and consequently has limited support for population genetics summary statistics. `PopGenome` (Pfeifer et al. 2014) offered a comprehensive R-based toolkit but has been removed from CRAN and is no longer maintained. `scikit-allel` (Miles et al. 2023) brought population genetic analyses into the Python ecosystem with NumPy-based array operations, offering a broad repertoire of statistics including haplotype-based selection scans and *F*-statistics; however, it is no longer under active development. `pixy` (Korunes and Samuk 2021; Bailey, Stevison, and Samuk 2025) addressed the important biases that missing data introduce in estimates in *π, d*_*XY*_, *F*_*ST*_, Tajima’s *D*, and *θ*_*W*_, at the cost of computational performance. `clam` is a very recent tool that focuses on faster computation of three statistics (*π, d*_*XY*_, and *F*_*ST*_) while accounting for missingness due to differences in callability across samples (Mirchandani et al. 2025). Lastly, `tskit` (Jeffery et al. 2026) provides exceptionally effcient computation of a general class of statistics, which is enabled by a completely different data format, the tree sequence, rather than the traditional genotype matrix (for example, in `VCF` or `PLINK` formats). However, the tree sequences need to be first inferred from the genotype data itself, a process which may not be feasible for some datasets, for example due to the lack of phased haplotypes.

Computation of population genetic statistics across all of these earlier tools is performed on the CPU, and statistics are typically computed sequentially along genomic windows. For large-scale projects involving windowed genome scans of tens of statistics across many populations, wall-clock times of hours to days are common. This limitation is particularly acute for two-locus statistics such as LD, where pairwise computations over variant sites are inherently *O*(*n*^2^) and form a bottleneck in inference pipelines such as `moments` (Jouganous et al. 2017; Ragsdale and Gravel 2019). Table 1 summarizes the coverage and capabilities of the most commonly used CPU tools alongside `pg_gpu` (this work).

**Table 1:** Coverage of population-genetics summary statistics across established tools that operate on a genotype matrix (VCF/zarr) and pg_gpu. ✓, full support; ~, partial or restricted support; —, no support or out of scope. Statistic categories follow the broad families described in the text. `tskit` (Jeffery et al. 2026) is omitted because it operates on tree sequences rather than a genotype matrix; `pg_gpu` accepts both via `HaplotypeMatrix.from ts` (right-most column). `moments` (Jouganous et al. 2017) is omitted because its scope is downstream demographic inference rather than genome-scale summary-statistic computation; `pg_gpu` provides a drop-in replacement for its LD-statistics parsing module (Supplementary Figure S7).

| Tool | Backend | Diversity<br>& SFS | Divergence<br>& *F*-stats | LD | Selection<br>scans | PCA | Local PCA<br>/ lostruct | GPU | Tree-seq.<br>input |
| --- | --- | --- | --- | --- | --- | --- | --- | --- | --- |
| VCFtools (Danecek et al. 2011) | C++ | ~ | ~ | ~ | — | — | — | — | — |
| PopGenome <sup>a</sup> (Pfeifer et al. 2014) | R | ✓ | ✓ | ~ | ~ | — | — | — | — |
| PLINK2 (Chang et al. 2015) | C++ | ~ | ~ | ~ | — | ✓ | — | — | — |
| ANGSD (Korneliussen, Albrechtsen, and Nielsen 2014) | C++ | ✓ | ✓ | — | — | ~ | — | — | — |
| scikit-allel <sup>b</sup> (Miles et al. 2023) | Python | ✓ | ✓ | ✓ | ✓ | ✓ | — | — | — |
| pixy (Korunes and Samuk 2021) | Python | ~ | ~ | — | — | — | — | — | — |
| clam (Mirchandani et al. 2025) | Python | ~ | ~ | — | — | — | — | — | — |
| pg-gpu (this work) | Python / CUDA | ✓ | ✓ | ✓ | ✓ | ✓ | ✓ | ✓ | ✓ |

<sup>a</sup>Removed from CRAN; no longer maintained. <sup>b</sup>Active maintenance has ceased.

Modern graphics processing units (GPUs) offer massive parallelism that is well-suited to the data-parallel structure of population genetics computation. Many statistics reduce to operations over matrices of haplotypes or genotypes that can be decomposed into independent per-site or per-window calculations, making them natural candidates for GPU acceleration. While GPUs have been applied to specific problems in population genetics—notably forward-in-time simulation (Lawrie 2017) and demographic inference via diffusion approximations (Gutenkunst 2021) or gradient-based optimization (Terhorst 2025) —no general-purpose GPU-accelerated library for computing population genetics summary statistics from empirical data has been developed.

Here, we present `pg_gpu`, a Python library that leverages NVIDIA GPUs via `CuPy` (Okuta et al. 2017) to compute a comprehensive catalog of population genetics statistics. `pg_gpu` implements over 70 functions spanning diversity, divergence, the site frequency spectrum, linkage disequilibrium, haplotype-based selection scans, admixture statistics, PCA, and relatedness—covering the large majority of statistics in routine use. The library uses fused CUDA kernels — kernels that compute many related statistics in a single pass over the data — and implements the generalized *θ*-estimation framework of Achaz (2009) to effciently derive all standard diversity and neutrality test statistics from a single site frequency spectrum computation. `pg_gpu` also provides a drop-in replacement for the LD-statistics parsing module of `moments`, achieving roughly 1,750-fold speedups while maintaining numerical accuracy at machine precision. Across windowed genome scans, `pg_gpu` delivers a median 139-fold and up to 1,096-fold speedup relative to `scikit-allel`. `pg_gpu` is open source and freely available at https://github.com/kr-colab/pg_gpu.

### Design and Implementation

`pg_gpu` is a pure-Python library built on `CuPy` (Okuta et al. 2017), with around 50 custom CUDA kernels (`cupy.RawKernel`) and dense linear algebra dispatched to `cuBLAS`; it depends only on `CuPy`, `NumPy` (Harris et al. 2020), `SciPy` (Virtanen et al. 2020), `scikit-allel` (Miles et al. 2023), `msprime` (Baumdicker et al. 2022), and `zarr`, and is distributed as a `pixi` environment so the full analysis stack installs with one command. Two thin wrapper classes—`HaplotypeMatrix` for phased haplotypes and `GenotypeMatrix` for unphased diploid genotypes—carry data through every analysis path; either can be built from a tree sequence (Jeffery et al. 2026), a `zarr` store (`bio2zarr`/`VCZ` (Czech et al. 2024) or `scikit-allel` layouts), or an in-memory `NumPy` array, and moves between CPU and GPU backings transparently.

The performance of `pg_gpu` rests on a single observation: most population-genetics statistics are per-site or per-pair reductions that spend most of their CPU time reading the genotype matrix from memory rather than doing arithmetic. `pg_gpu` therefore uses *fused* kernels: one kernel reads each entry of the matrix once and computes whichever statistics from a related family the user asked for. A single entry point, `windowed analysis`, takes a list of statistics and runs the fewest such kernels that together cover the request. If the data are too large to fit in GPU memory at once, the same kernels stream over the input in chunks whose size is picked automatically, so a whole chromosome arm of ~10 million variants fits on one A100 without any tuning by the user. Full details—the array layouts, the fused-kernel families, how missing data and accessibility masks are handled, the chunking rule, and the downstream interfaces—are described in the Supplementary Material (Extended Design and Implementation).

### Features

`pg_gpu` provides a comprehensive catalog of population-genetics statistics organized into eleven categories (Table 2): classic estimators (*π, θ*_*W*_, Tajima’s *D*, Hudson and Weir– Cockerham *F*_*ST*_, *d*_*XY*_, the unfolded and joint SFS, Patterson’s *f*_2_/*f*_3_/*D*, iHS, nSL, Garud’s *H*) alongside several less commonly supported additions useful in modern workflows—the Schrider et al. (Schrider et al. 2018) distance-based two-population statistics, the RAiSD *µ* components (Alachiotis and Pavlidis 2018), the diplotype-frequency spectrum, and a `FrequencySpectrum` interface to the Achaz (Achaz 2009) weighted-SFS framework for custom *θ* estimators. Almost every statistic is also reachable through `windowed analysis`, the entry point for genome scans using the fused kernels (Design and Implementation). The same entry point applies whether the input is an in-memory matrix or a chunked `VCZ` store streamed across the chromosome. A complete listing of all implemented statistics with primary references, together with an extended description of the API and resampling utilities, is provided in the Supplementary Material (Table S2).

**Table 2:**
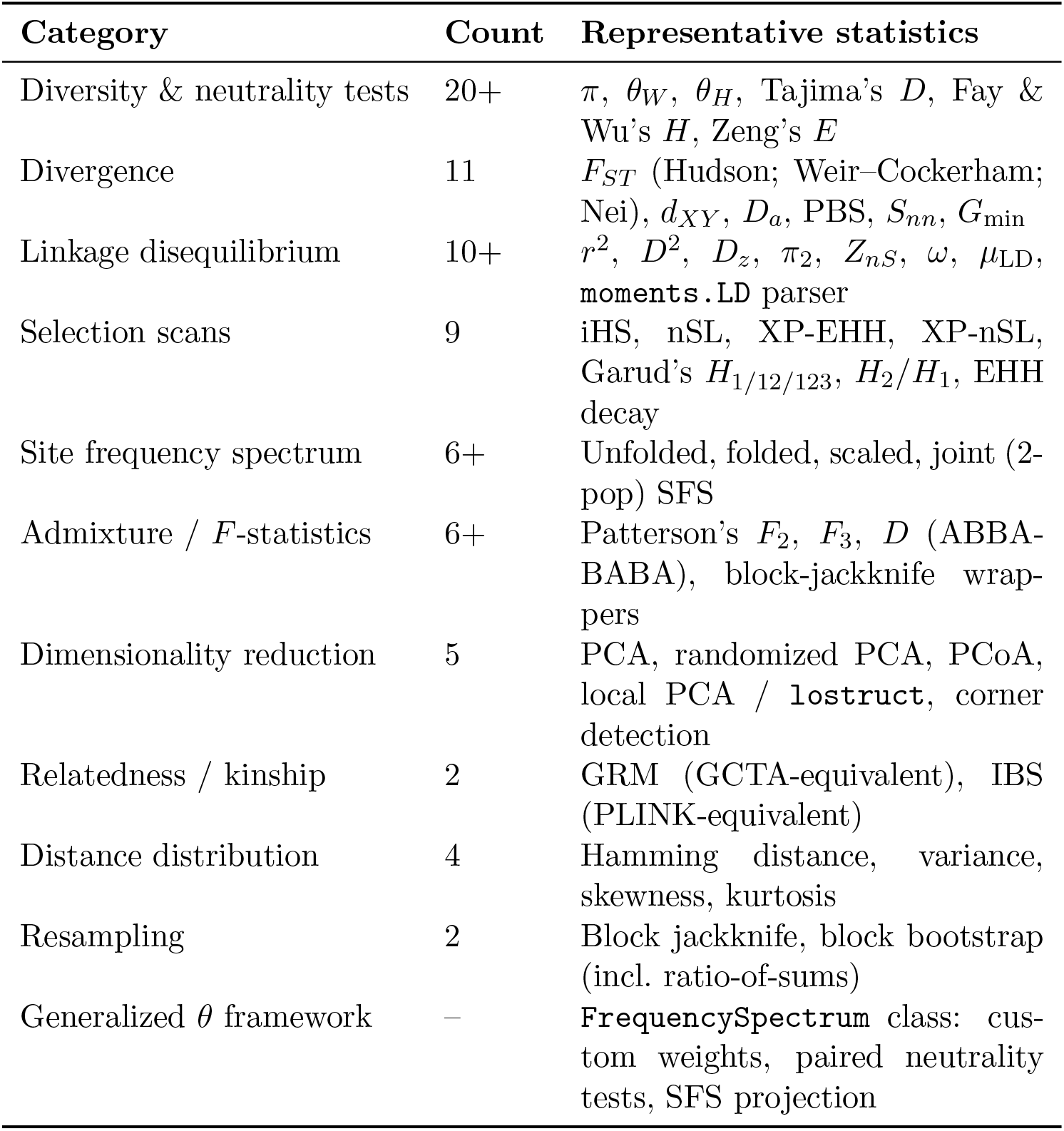
Summary of statistics implemented in pg_gpu. The full catalog with citations is provided in Supplementary Table S2.

| Category | Count | Representative statistics |
| --- | --- | --- |
| Diversity & neutrality tests | 20+ | $\pi$, $\theta_W$, $\theta_H$, Tajima’s $D$, Fay & Wu’s $H$, Zeng’s $E$ |
| Divergence | 11 | $F_{ST}$ (Hudson; Weir–Cockerham; Nei), $d_{XY}$, $D_a$, PBS, $S_{nn}$, $G_{\min}$ |
| Linkage disequilibrium | 10+ | $r^2$, $D^2$, $D_z$, $\pi_2$, $Z_{nS}$, $\omega$, $\mu_{LD}$, `moments`.LD parser |
| Selection scans | 9 | iHS, nSL, XP-EHH, XP-nSL, Garud’s $H_{1/12/123}$, $H_2/H_1$, EHH decay |
| Site frequency spectrum | 6+ | Unfolded, folded, scaled, joint (2-pop) SFS |
| Admixture / $F$-statistics | 6+ | Patterson’s $F_2$, $F_3$, $D$ (ABBA-BABA), block-jackknife wrappers |
| Dimensionality reduction | 5 | PCA, randomized PCA, PCoA, local PCA / `lostruct`, corner detection |
| Relatedness / kinship | 2 | GRM (GCTA-equivalent), IBS (PLINK-equivalent) |
| Distance distribution | 4 | Hamming distance, variance, skewness, kurtosis |
| Resampling | 2 | Block jackknife, block bootstrap (incl. ratio-of-sums) |
| Generalized $\theta$ framework | – | `FrequencySpectrum` class: custom weights, paired neutrality tests, SFS projection |

## Results

### Benchmark dataset

For benchmarking, we use Phase 3 of the *Anopheles gambiae* 1000 Genomes Project (Ag1000G) (Anopheles gambiae 1000 Genomes Consortium 2017), a deep-coverage whole-genome resequencing panel of 1,470 phased mosquitoes (2,940 haplotypes) sampled from 22 populations across sub-Saharan Africa. Chromosome arm 3R alone carries 10.9 million phased, biallelic SNPs after filtering and accessibility masking; the 53 Mb arm contains multiple known segregating inversions and well-characterized insecticide-resistance sweeps, and is among the most variable autosomal regions sequenced in any eukaryote. The combination of extreme levels of segregating variation, non-trivial sample size, natural missingness, and rich population structure makes it a stringent and biologically-meaningful stress-test for the fused CUDA kernels. Where a ground truth is needed—for missing-data bias, runtime scaling, and demographic-inference cross-checks against `moments`—we pair Ag1000G with `msprime` (Baumdicker et al. 2022) simulations so each result is supported by both empirical and simulation evidence.

### Accuracy, performance, and scaling

We first verified that `pg_gpu` reproduces results from established CPU implementations: every statistic with a counterpart in `scikit-allel`, `PLINK2`, or `moments` matches the reference to within floating-point error, with per-statistic comparisons reported in the Supplementary Material (Table S1, Supplementary Figures S5 and S9).

Across the 23 Ag1000G 3R statistics with a directly comparable `scikit-allel` implementation (2,940 haplotypes, 10.9 million variants), the median `pg_gpu` speedup is 139× and the maximum 1,096× (Weir–Cockerham *F*_*ST*_; Figure 1, Supplementary Figure S1): divergence and pairwise-LD kernels gain the most (600–1,100×), single-population diversity and SFS statistics 130–380×, and haplotype-based selection scans 9–56×, the last bounded by per-haplotype work that `scikit-allel`’s C extensions already vectorize. On a 50 kb-windowed slice of the same data, `pg_gpu` and `scikit-allel` estimates of *π*, an *R*_0_-style heterozygosity ratio, Tajima’s *D*, and LD decay overlay to within line width while the windowed scan and LD-decay computation run 5.8× and 323× faster (Supplementary Figure S3); a controlled cross-check on simulated two-population `msprime` data reproduces the same speedup pattern against both `scikit-allel` and `PLINK2` (Supplementary Figure S2).

**Figure 1:**
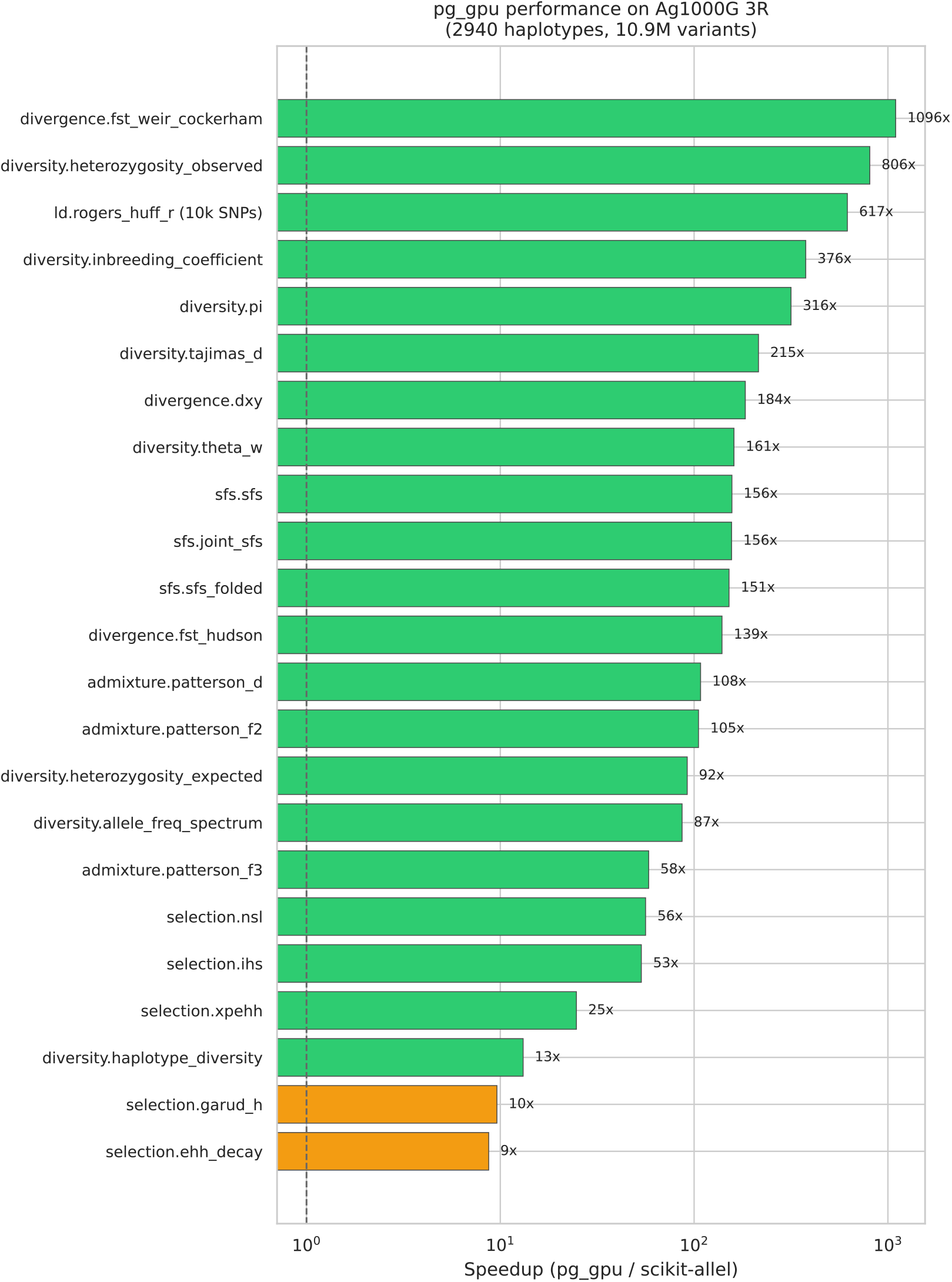
`pg_gpu` speedup over `scikit-allel` on Ag1000G Phase 3 chromosome 3R (2,940 haplotypes, 10.9M variants). The all-pairwise Rogers–Huff LD comparison is run on the first 10,000 SNPs (200 haplotypes) to keep the `scikit-allel` reference tractable; all other statistics are on the full chromosome arm. Bars show wall-clock speedup on a single NVIDIA A100 GPU; orange bars indicate statistics where `scikit-allel`’s existing optimized implementation limits the achievable gain (typically < 10×).

A separate bottleneck is the tallying of multi-population two-locus haplotypes into LD statistics used for demographic inference (Ragsdale and Gravel 2019). On a four-population coalescent simulation (105 LD statistics, 10 recombination-distance bins, 10 replicates of 1 Mb) `pg_gpu` agrees with `moments`-LD to a maximum relative error of 1.8 × 10^−11^ while cutting per-replicate parsing time from 256 s to 0.2 s—a 1,756 *±* 410× speedup (Supplementary Figure S7)—making whole-genome LD-based inference tractable on standard hardware; a per-population-count breakdown is given in Supplementary Figure S8.

Runtime scaling was characterized along two axes with `msprime` simulations (Baumdicker et al. 2022): sample size (100 to 100,000 haplotypes at a fixed 100,000 variants) and variant count (~ 10^3^ to ~ 10^6^ segregating sites at a fixed 200 haplotypes, with realistic LD; Supplementary Figure S4). Per-site statistics scale linearly on both axes and finish in well under a second across the tested range. Garud’s *H*, a haplotype-pattern statistic computed via per-window haplotype hashing, also scales linearly in sample size but with a larger per-sample constant. The two-locus statistic *Z*_*nS*_ (a sampled-pairs estimator) shows a sub-quadratic profile in sample size that plateaus once the sampled-pair budget saturates. At the largest configuration tested (100,000 haplotypes × 100,000 variants) every single-pass statistic completes in under ten seconds on one A100.

### Benchmarks and Applications

We illustrate two end-to-end uses of `pg_gpu` on data and methodology where the GPU implementation enables an analysis that is slow or intractable on CPU. A third use—supplying `pg_gpu`-computed LD statistics to the `moments` demographic-inference workflow as a single-import drop-in replacement, with downstream parameter estimates indistinguishable from an analysis using the native parser—is described in the Supplementary Material (Supplementary Figure S12).

#### Benchmarking a whole-arm Ag1000G genome scan

Partitioning the 1,470 Ag1000G samples into West and East African groups (200 diploids / 400 phased haplotypes each) and running a full population-genetic scan of chromosome 3R—loading 10.9 million phased variants from `zarr`, attaching the canonical accessibility mask, transferring to GPU, and computing 19 windowed statistics in 100 kb / 10 kb windows plus per-population and joint SFS—takes 168 seconds total wall time, of which only 30 seconds is GPU compute (Figure 2). The same workflow expressed in `scikit-allel` runs in roughly 75 minutes (windowed statistics alone), so a single A100 reduces the runtime from over an hour to seconds while recovering the expected genome-wide signatures: *π* heterogeneity, negative Tajima’s *D* consistent with non-equilibrium demography and selection, and elevated Hudson *F*_*ST*_ between subspecies. Every per-window panel comes from a single call to `pg_gpu`’s `windowed_analysis` function, which will calculate multiple statistics (e.g., this plot) from the loaded haplotype matrix in a single forward pass.

**Figure 2:**
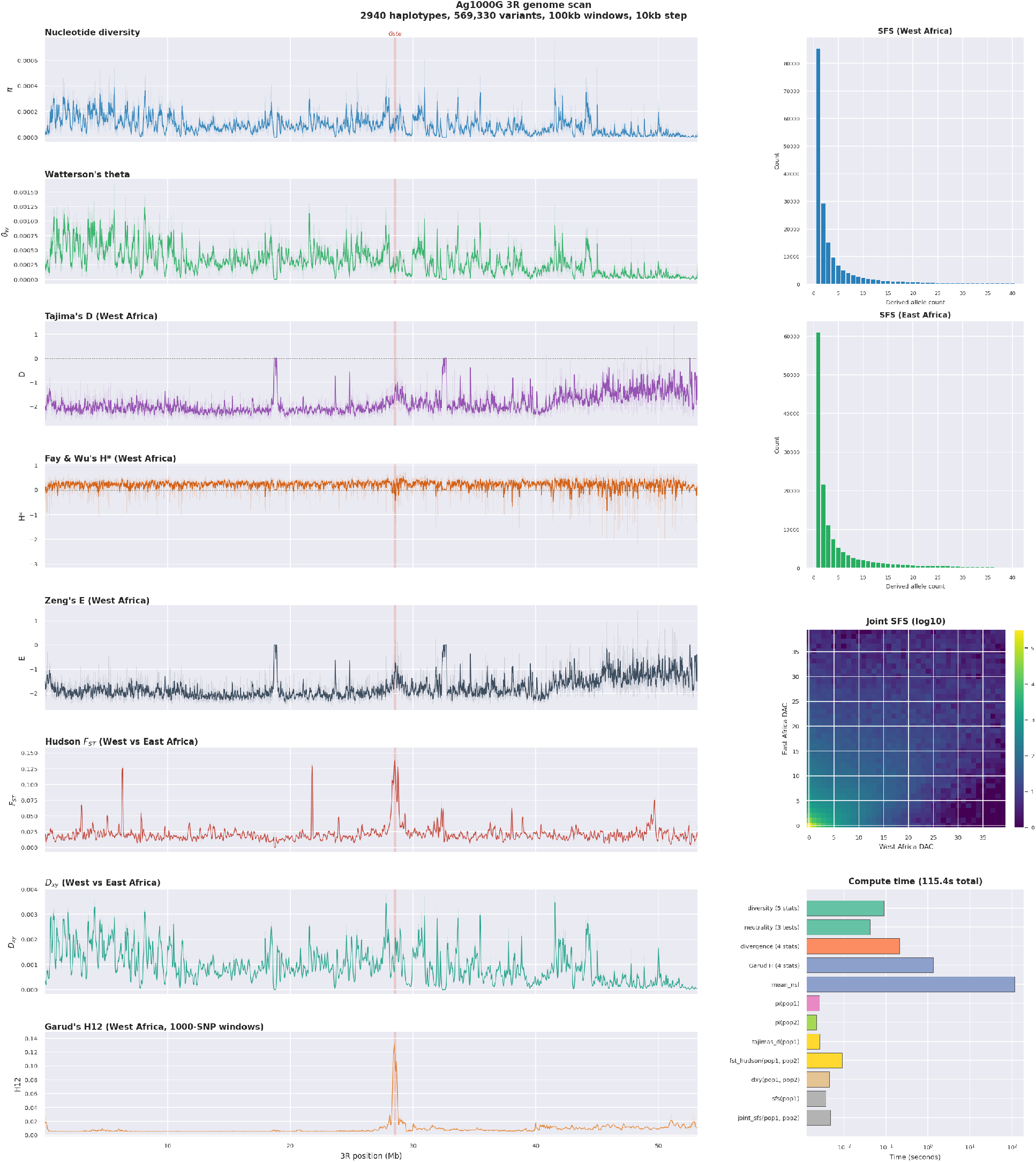
Multi-statistic scan of Ag1000G chromosome 3R using `pg_gpu`. Left column, eight per-window panels in 100 kb windows with a 10 kb step (*H*_12_ uses 1,000-SNP windows on its own grid): nucleotide diversity *π*, Watterson’s *θ*_*W*_, Tajima’s *D*, Fay & Wu’s *H*^∗^, and Zeng’s *E* (all West Africa); Hudson *F*_*ST*_ and *d*_*XY*_ between West and East Africa; and Garud’s *H*_12_ (West Africa). The red band on every left-column panel marks the *Gste* locus (3R:28.48–28.60 Mb), a well-characterised insecticide-resistance locus that the scan recovers as coordinated drops in *π* and *θ*_*W*_ with elevated *H*_12_. Right column: West African unfolded SFS, joint SFS heatmap (West vs. East), and per-step compute time. Total compute time is 30 s on a single A100 GPU; total wall time including I/O is 168 s.

#### Whole-genome local PCA

The local-PCA / `lostruct` (H. Li and Ralph 2019) analysis workflow—comprised of per-window local PCA, MDS using Frobenius distance between window covariances, and corner detection—identifies genomic intervals whose structure deviates from the genome-wide pattern. As a sanity check with a controlled signal we applied it to a coalescent simulation (Baumdicker et al. 2022) containing a single hard sweep (Supplementary Figure S10): the embedding cleanly separates neutral, linked, and sweep windows and the leading MDS axis tracks the localized peak of Garud’s *H*_12_ at the sweep focal site. Applied to the West African subset of Ag1000G 3R (100 diploids / 200 phased haplotypes, 10.9M variants, ~10,900 non-overlapping 1,000-SNP windows; Figure 3), three distinct corner clusters are recovered, and the corner-1 region near ~28 Mb overlaps with a localized peak in the companion Garud’s *H*_12_ track (the *Cyp6* /*Gste* insecticide-resistance locus on 3R), a known sweep target in West African *An. gambiae*.

**Figure 3:**
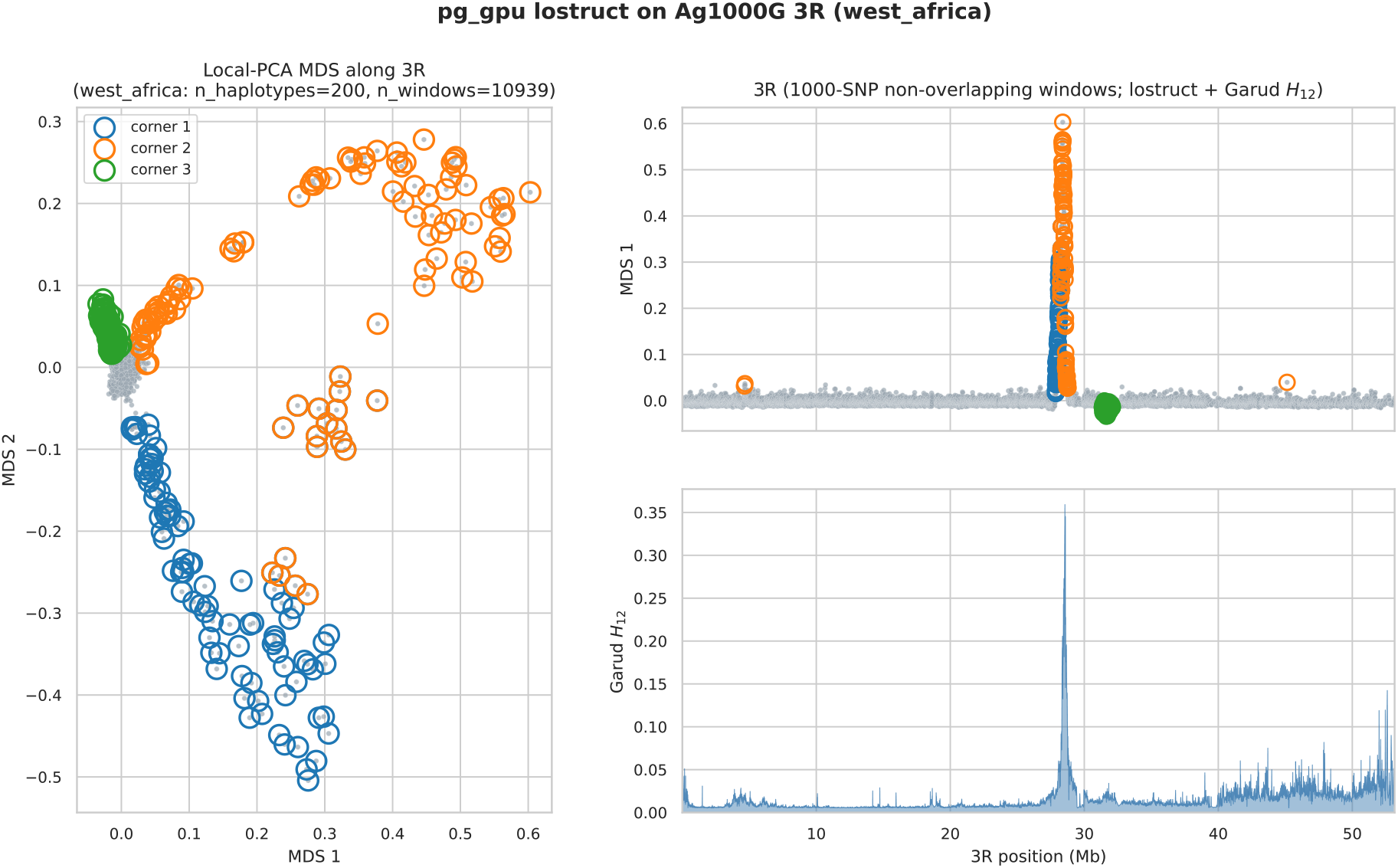
`lostruct` on the West African subset of Ag1000G 3R (100 diploids / 200 phased haplotypes, 10.9M variants, ~10,900 non-overlapping 1,000-SNP windows). Left: MDS embedding of windowed PCA distances; windows in each of three detected corner clusters are circled. Right: `lostruct` MDS axis 1 along the chromosome (top) and companion Garud’s *H*_12_ track on the same windows (bottom).

#### Streaming a biobank-scale genome scan

At biobank sample sizes (e.g. ≥ 10^5^ individuals) a single chromosome will not fit on a GPU all at once. `pg_gpu` handles this by reading the chromosome one chunk at a time; the same windowed-statistic, SFS, LD, and pairwise-relatedness functions used on genotype matrices that can fit in memory work without change on the chunked input. We demonstrate this on chromosome 15 of `stdpopsim`’s `OutOfAfrica 2T12` model simulated at 50,000 diploid individuals per population (200,000 haplotypes, 11.6 million variants, ~ 2.3 TB on disk as an uncompressed text VCF). The full scan shown in Supplementary Figure S11—windowed diversity and divergence across all 200,000 haplotypes at three window sizes, marginal SFS per population, a joint SFS projected from the full panel to a smaller grid via per-variant hypergeometric sampling, plus Garud’s *H* and an LD decay curve on smaller per-population subsamples—finishes in about 16 minutes on a single A100 80 GB. GPU memory use stays around 12 GB regardless of how long the chromosome is.

## Discussion

`pg_gpu` fills a long-standing gap in the population-genetics software ecosystem. Existing tools optimize specific axes—`VCFtools` and `PLINK` for command-line access to focused sets of statistics, `scikit-allel` for a Python-native research workflow, `pixy` for unbiased estimation under missingness, and `tskit` for tree-sequence inputs, but none brings the full catalog of routine summary statistics onto modern GPU hardware. By unifying eleven categories of statistics under a single Python API on top of `CuPy`, `pg_gpu` turns analyses that were previously hours-long serial CPU scans (e.g. full chromosome arm scans using `scikit-allel`) or computationally infeasible (chromosome-scale LD calculation using all SNP pairs) into runs of seconds to minutes on a single GPU.

Three design choices that emerged from this work may be useful to other authors of GPU-accelerated scientific software. First, fused kernels that emit any subset of a related family of statistics from registers in a single launch substantially outperform sequential per-statistic GPU calls, because the dominant cost on this class of workload is reading the haplotype matrix off-RAM rather than the arithmetic itself. Second, memory-aware chunking lets the same kernels run on data far larger than GPU memory without user tuning. The route is not new: chunked, columnar formats like `VCZ` (Czech et al. 2024) are the standard way to store chromosome-scale variant data. `VCZ` stores the genotype matrix as a grid of rectangular blocks (variants by samples), with each block compressed independently, so any region of the genome can be reconstructed by reading just the blocks that overlap it. `pg_gpu` reads those blocks one at a time and feeds each to the same fused kernels used on in-memory data, accumulating per-window results across the chromosome. A crucial aspect of our chunking strategy is to leverage `kvikio` (RAPIDS Development Team 2021) together with NVIDIA’s nvCOMP GPU decompressors so that each compressed `VCZ call genotype` chunk is decoded directly on the GPU. The potential of `kvikio`’s GPU-native `zarr` backend for accelerating the processing of population genetic data was pointed out by Czech et al. (2024), and has proven essential for the effciency of the streaming-to-GPU strategy implemented in `pg_gpu`. Because the per-site and per-window summaries each kernel emits compose across blocks, the streamed result matches the in-memory result; chunking is a strategy bound memory usage, not an approximation. Third, processing windows in tiles—so that working memory scales with the tile size 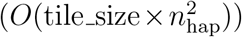 rather than with the number of windows 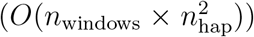—lets the same code path scale from a single chromosome to a whole genome.

Several limitations are worth stating. `pg_gpu` requires an NVIDIA GPU because `CuPy` targets CUDA; ports to AMD ROCm or Apple Metal would require non-trivial kernel rewrites. Haplotype-pattern statistics that scale as 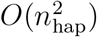 in sample size remain bound by GPU memory at large scales (we tested up to 100,000 haplotypes); genuine biobank-scale sample-sizes will need additional chunked-pair kernels for the haplotype-pair statistics. The randomized SVD used for the local PCA is an approximation to the decomposition of the full covariance matrix; although a dense eigendecomposition is available when exact `lostruct` outputs are required. Finally, `pg_gpu` currently supports phased haplotypes and unphased diploid genotypes but does not yet handle low-coverage genotype likelihoods natively (cf. `ANGSD`); extending the framework to genotype likelihoods is straightforward in principle and is part of planned future work.

Looking forward, we see `pg_gpu` as a foundation for two further directions: integration with simulation-based machine-learning frameworks for demographic and selection inference, where rapid summary-statistic computation is the bottleneck during training; and as a reference implementation of the Achaz (Achaz 2009) weighted-SFS framework, lowering the barrier for population geneticists to design and validate custom test statistics. We expect both will broaden the set of analyses that fit comfortably within standard whole-genome research workflows.

## Availability

`pg_gpu` is freely available at https://github.com/kr-colab/pg_gpu.

## Supplementary Information

An extended description of the design and implementation, the full statistic catalog with primary references, an extended description of the API and features, numerical accuracy comparisons against `scikit-allel`, `PLINK2`, and `moments`, a worked `moments`-LD demographic-inference example, and supplementary figures (including a breakdown of the speed and accuracy of LD statistics parsing for `moments`) are provided in the Supplementary Material.

## Acknowledgments

We thank members of the Kern-Ralph co-lab for feedback on the library and the manuscript.

## Funding

This work was supported by NIH grants R01HG010774, R01HG012473, and R35GM148253.

## Generative AI use

Anthropic’s Claude (Opus 4.6) was used to assist in drafting and optimizing the fused `cupy.RawKernel` CUDA implementations underlying `pg_gpu`. All model-generated kernel code was reviewed, profiled, and validated by the authors against appropriate reference implementations (Supplementary Table S1) before inclusion in the final codebase.

## Notes

https://github.com/kr-colab/pg_gpu/

### Competing Interest Statement

The authors have declared no competing interest.

