## Supplemental Figures for "GPU accelerated population genetics statistics using pg_gpu"

### Supplementary Material

#### Numerical accuracy

We validated `pg_gpu` against `scikit-allel` using a 4 Mb region of Ag1000G chromosome 3L (3L:1,000,000–5,000,000; 2,940 haplotypes after removing missing-data sites). Statistics whose estimators reduce to exact integer counts—site-frequency spectra, two-locus haplotype-frequency tallies (Garud's $H$), and Patterson's $f_2$—agree exactly. Statistics that involve harmonic sums or unbiased corrections ($\theta_W$, Tajima's $D$) agree to a relative error of at most $5 \times 10^{-4}$ (Table S1). A directly comparable Hudson $F_{ST}$ computation against `PLINK2` on simulated `msprime` data agrees to $1.7 \times 10^{-8}$, and the corresponding PCA scores agree to $|r| > 0.88$ through PC5 (Figure S5). The two-population LD-decay curves estimated by `pg_gpu` and `moments` on identical replicates are likewise numerically indistinguishable across all nine LD statistics shown (Figure S9).

Table S1: Agreement between `pg_gpu` and `scikit-allel` on Ag1000G 3L (2,940 haplotypes, 436,555 biallelic sites). Scalar statistics report the relative error $|x_{\text{pg}} - x_{\text{al}}|/|x_{\text{al}}|$. Vector statistics (SFS, windowed scans) report the Pearson correlation across $n$ entries; for windowed statistics the median per-window relative error is also given.

| **Category** | **Statistic** | pg_gpu | `scikit-allel` | **Rel. error** |
|---|---|---|---|---|
| Diversity | $\pi$ | $1.8660 \times 10^{-3}$ | $1.8660 \times 10^{-3}$ | $2.5 \times 10^{-7}$ |
| | $\theta_W$ | $3.2257 \times 10^{-3}$ | $3.2250 \times 10^{-3}$ | $2.1 \times 10^{-4}$ |
| | Tajima's $D$ | $-1.3642$ | $-1.3635$ | $5.1 \times 10^{-4}$ |
| Divergence | Hudson $F_{ST}$ | $2.9060 \times 10^{-3}$ | $2.9060 \times 10^{-3}$ | $2.8 \times 10^{-15}$ |
| | Weir–Cockerham $F_{ST}$ | $2.9262 \times 10^{-3}$ | $2.9262 \times 10^{-3}$ | $1.0 \times 10^{-15}$ |
| | $d_{XY}$ | $1.9002 \times 10^{-3}$ | $1.9002 \times 10^{-3}$ | $1.1 \times 10^{-16}$ |
| Selection | Garud's $H_1$ | $5.000 \times 10^{-3}$ | $5.000 \times 10^{-3}$ | 0 |
| | Garud's $H_{12}$ | $5.050 \times 10^{-3}$ | $5.050 \times 10^{-3}$ | 0 |
| Admixture | Patterson's $F_2$ (mean) | $2.278 \times 10^{-4}$ | $2.278 \times 10^{-4}$ | 0 |
| SFS (Pearson $r$) | Unfolded SFS, $n$=201 | — | — | $r = 1.0000$ |
| | Joint SFS, $n$=5,174 | — | — | $r = 1.0000$ |
| Windowed (50 kb, $n$=80) | $\pi$ | — | — | $4.5 \times 10^{-15}$ |
| | $\theta_W$ | — | — | $1.6 \times 10^{-4}$ |
| | Hudson $F_{ST}$ | — | — | $1.9 \times 10^{-15}$ |

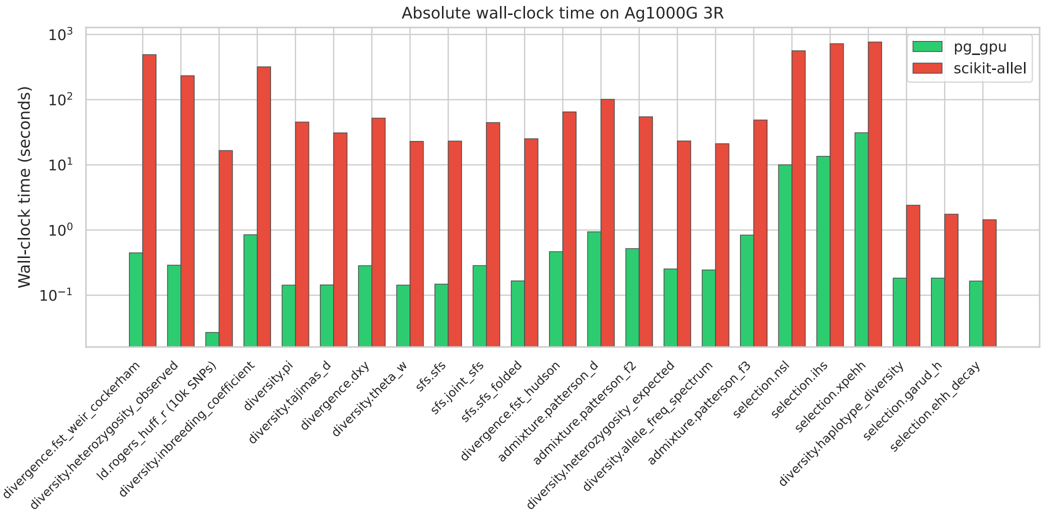


Figure S1: Absolute wall-clock time on Ag1000G chromosome 3R for the 23 statistics with both `pg_gpu` and `scikit-allel` implementations. Companion to Figure 1.

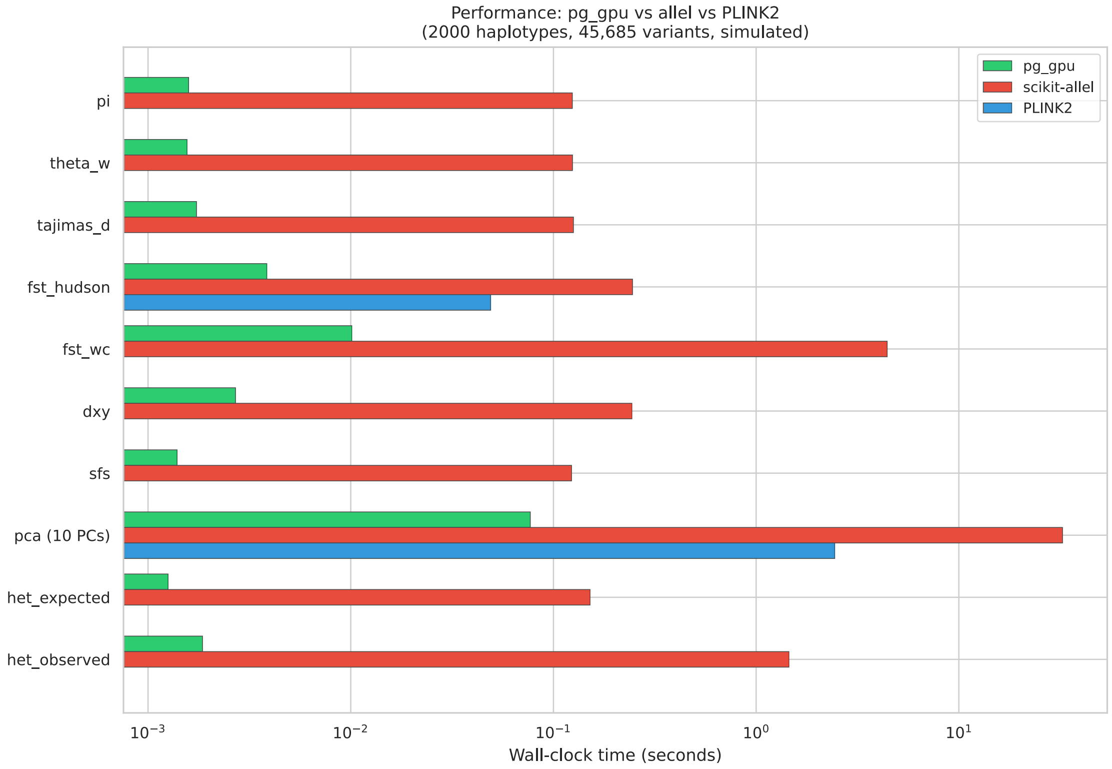


Figure S2: Performance comparison on simulated `msprime` data (two-population split, $N_e = 10^4$, 10 Mb sequence, 500 diploid individuals per population) against `scikit-allel` and `PLINK2` on shared statistics. The simulated benchmark provides a controlled cross-check of the Ag1000G results in Figure 1.

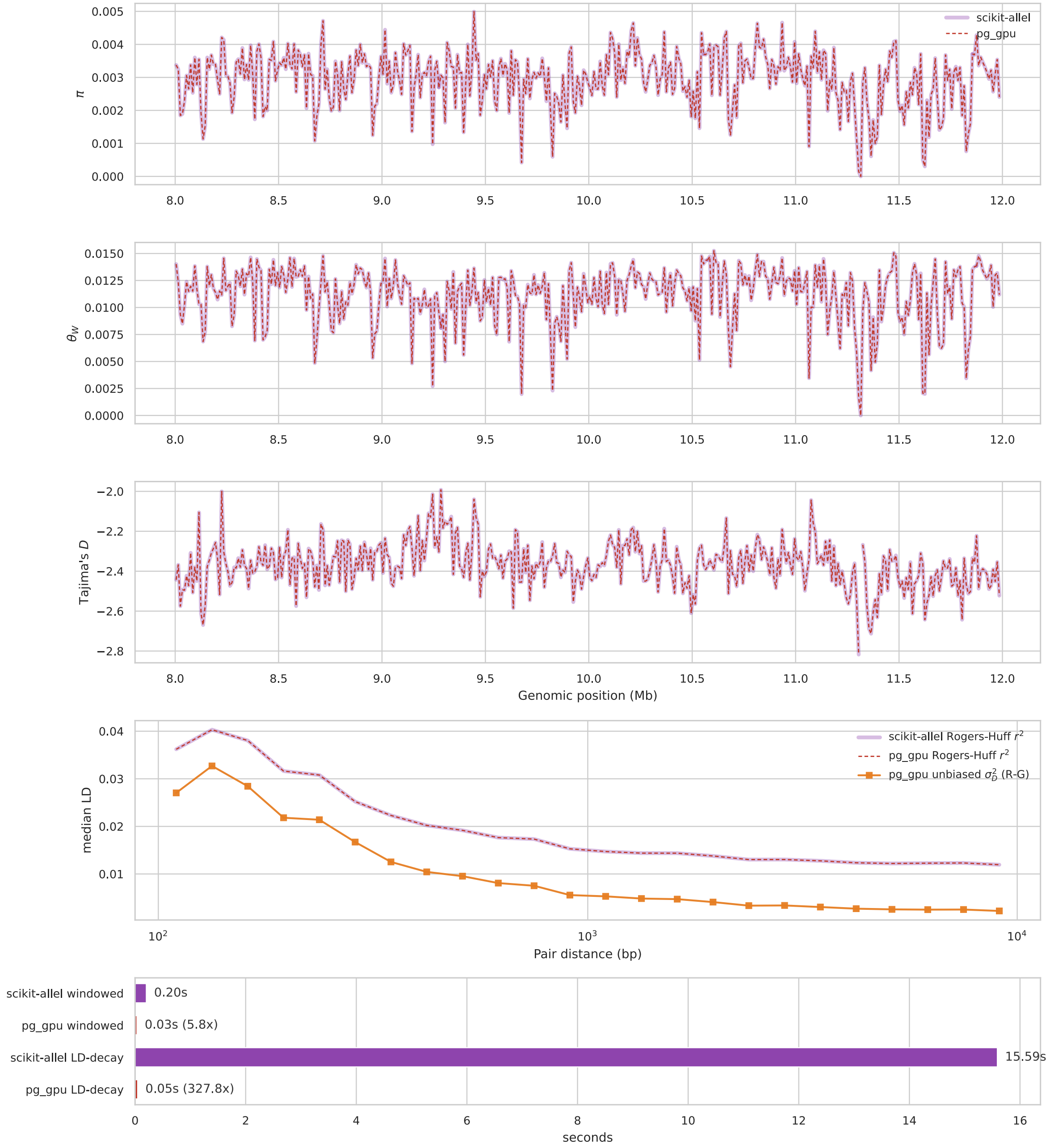


Figure S3: Equivalence and speedup of `pg_gpu` vs. `scikit-allel` on a single chromosome of phased Ag1000G data. Top three panels: nucleotide diversity ($\pi$), $R_0$, and Tajima's $D$ in 50 kb windows along the chromosome, computed by both tools—curves overlay to within line width. Lower panel: LD-decay (mean $r^2$ vs pairwise distance) showing identical estimates from `scikit-allel` and the two `pg_gpu` LD estimators (Rogers–Huff and unbiased $\sigma_d^2$). Horizontal bars at the bottom: wall-clock times for the windowed scan (5.8× faster) and LD-decay computation (323× faster).

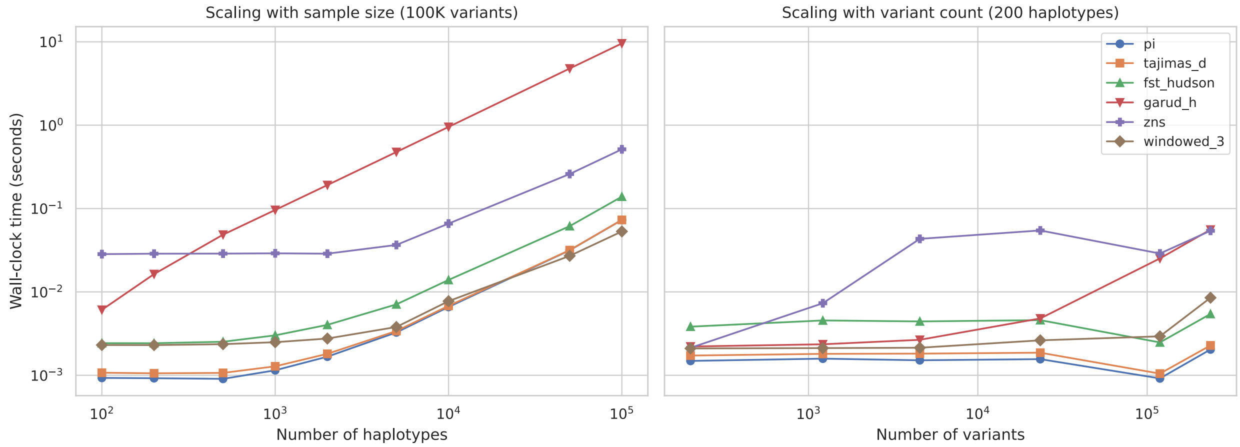


Figure S4: Runtime scaling with sample size (left, fixed at 100,000 variants) and variant count (right, fixed at 200 haplotypes). Both axes are log-scaled. `windowed_3` is a fused windowed kernel that computes $\pi$, $\theta_W$, and Tajima's $D$ in a single pass.

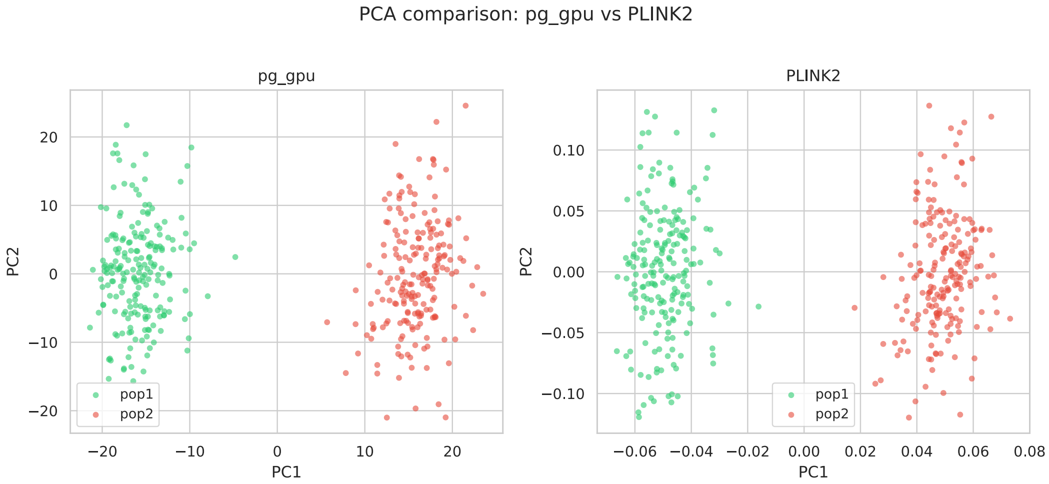


Figure S5: Per-axis correspondence of `pg_gpu` PCA scores against `PLINK2` on simulated data (800 haplotypes, 19,470 biallelic variants, two-population split). PC1 agrees to $|r| = 0.9997$; subsequent axes show progressively reduced absolute correlation that reflects the expected sensitivity of low-eigenvalue eigenvectors to floating-point roundoff, with $|r| > 0.88$ through PC5.

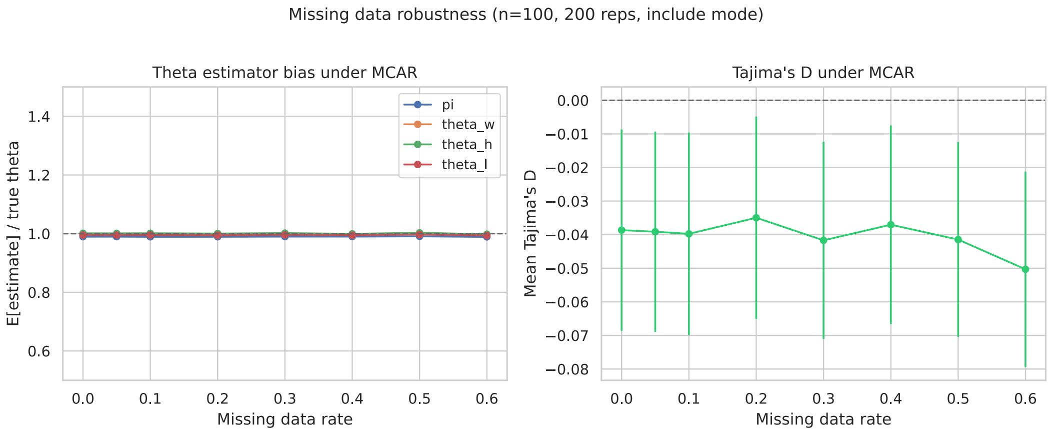


Figure S6: Bias of pg_gpu's nucleotide-diversity estimator under randomly induced missing genotypes (200 msprime replicates per missing-data rate). The estimator is unbiased across missing rates from 0 to 60%, mirroring the per-site valid-sample-count handling implemented for all single-population diversity estimators.

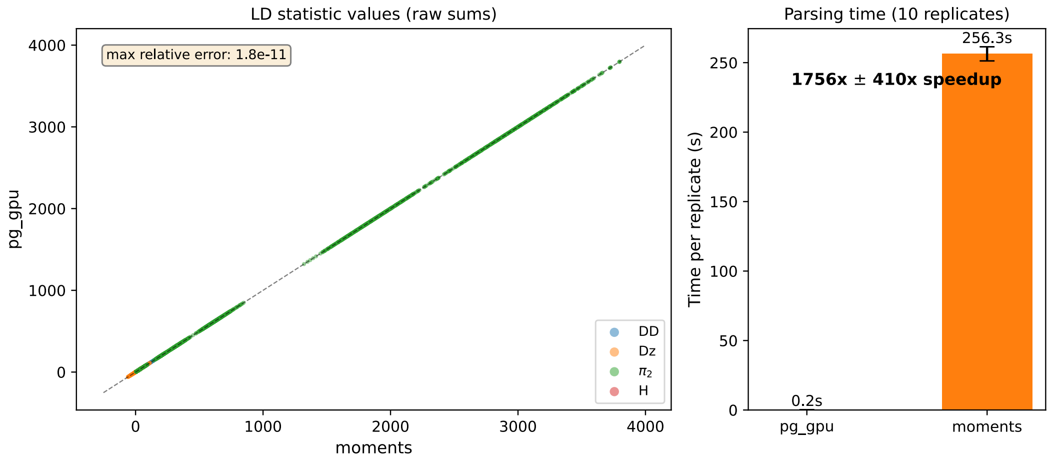


Figure S7: pg_gpu The accuracy and speed of parsing LD statistics, against moments. Left: raw two-locus LD-statistic sums ($D^2$, $Dz$, $\pi_2$, $H$) for a four-population split simulation (10 replicates; 105 statistics). Right: per-replicate parsing time on moments CPU vs. pg_gpu GPU ($1{,}756 \pm 410\times$ speedup).

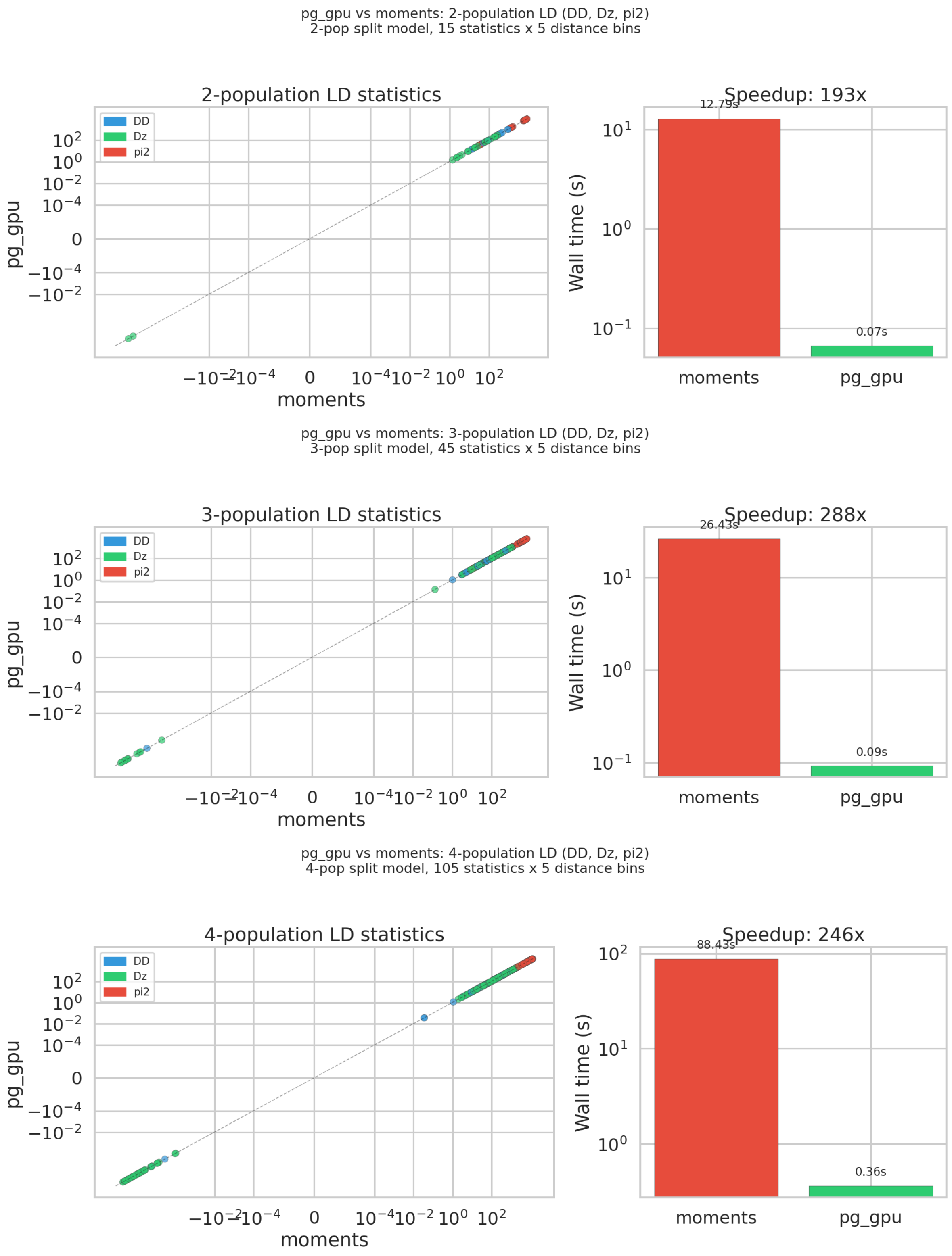


Figure S8: Numerical validation of 2-, 3-, and 4-population LD statistics on a 500 kb bp-binned stress test ($\mu = 10^{-7}$, no recombination map). `pg_gpu` agrees with `moments` to machine precision; reported speedups (193×, 288×, 246×) are lower than Figure S7 because that benchmark uses the slower r-binned `moments` path on 20× more sequence.

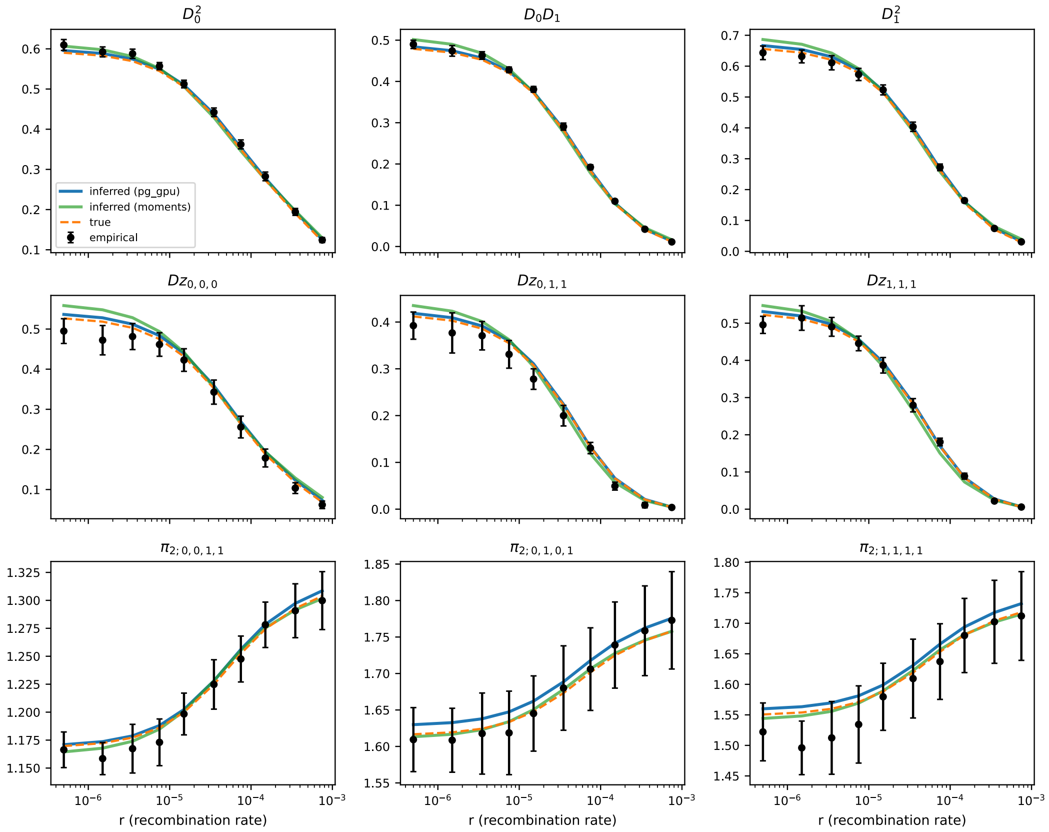


Figure S9: Two-population LD decay curves estimated by `pg_gpu` (this work) and `moments` on identical replicate simulations. The two parsers produce numerically identical outputs across all 9 LD statistics shown, confirming `pg_gpu` can serve as a transparent drop-in replacement for the native `moments.LD` parsing module.

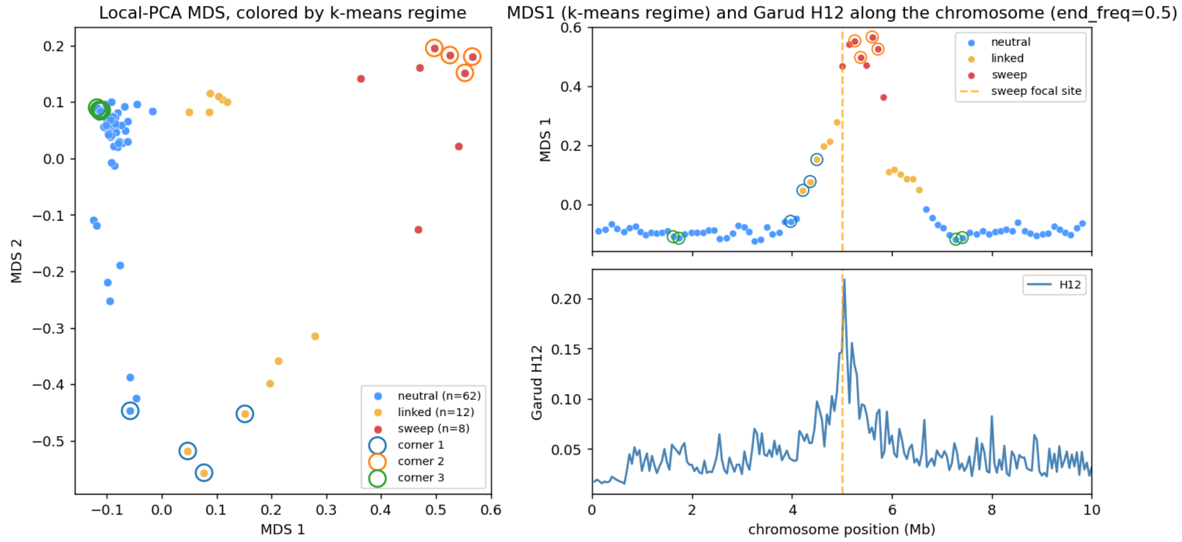


Figure S10: Local PCA (`lostruct`) on a coalescent simulation containing a hard sweep at 5 Mb. Left: MDS embedding of windowed PCA distances, colored by k-means regime (neutral, linked, sweep). Right: MDS axis 1 (top) and Garud's $H_{12}$ (bottom) along the chromosome.

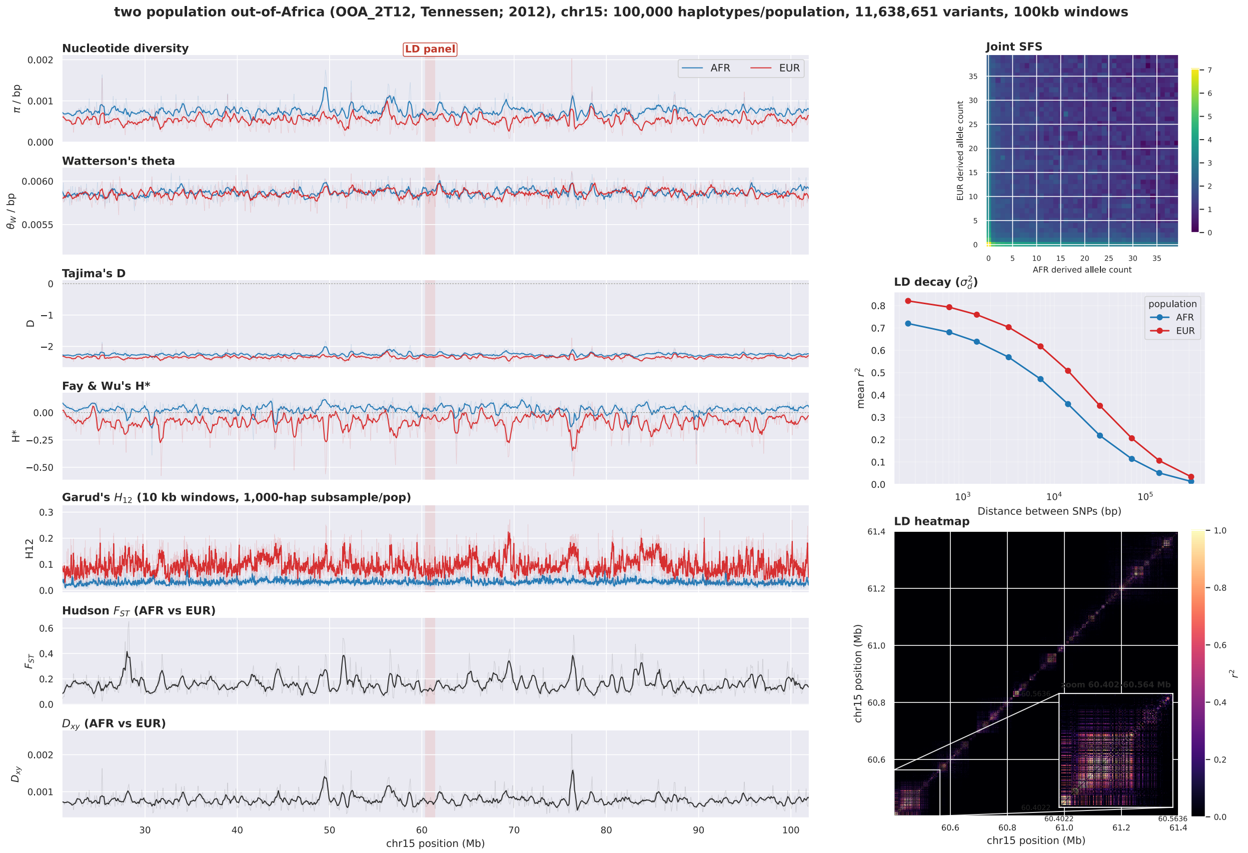


Figure S11: Biobank-scale genome scan of chromosome 15 from `stdpopsim`'s `OutOfAfrica_2T12` model, simulated at 50,000 diploids per population (200,000 haplotypes, 11.6 million variants). Left column shows the per-window scan across all 100,000 haplotypes per population: nucleotide diversity $\pi$, Watterson's $\theta_W$, Tajima's $D$, and normalized Fay & Wu's $H^*$ in 100 kb windows; Hudson $F_{ST}$ and $d_{XY}$ between populations; and Garud's $H_{12}$ in 10 kb windows on a 1,000-haplotype subsample per population (the Garud kernel caps near 1,024 haplotypes). The red band marks the 1 Mb sub-region used for the pairwise-$r^2$ heatmap. Right column, top to bottom: joint SFS projected from the full 100,000-haplotype panel per population to a 200-by-200 display grid via per-variant hypergeometric sampling (the full 100,000-by-100,000 histogram is both unviewable and larger than the GPU, but every variant from every haplotype contributes to the projected version); mean $r^2$ vs. distance between SNPs on common variants (minor-allele frequency $\geq 0.15$, 5,000-haplotype subsample per population) pooled across 16 probe regions tiling the chromosome; and the pairwise-$r^2$ heatmap of the 1 Mb sub-region (5,000-haplotype subsample, single population) with a zoom inset on the densest LD block. The complete scan finishes in about 16 minutes on a single A100 80 GB; GPU memory use stays around 12 GB regardless of chromosome length.

#### Demographic inference with pg_gpu LD statistics

Because pg_gpu's compute_ld_statistics returns LD-statistic sums in moments' canonical schema (Design and Implementation), it can replace the native moments.LD parser in an existing inference pipeline by changing a single import. We verified this end to end on a three-population isolation-with-migration model: fitting the model with pg_gpu-calculated LD statistics inside the standard moments workflow recovers parameter estimates indistinguishable from those obtained with the native parser (Figure S12), while computing the input statistics at the speedups reported in Figure S7.

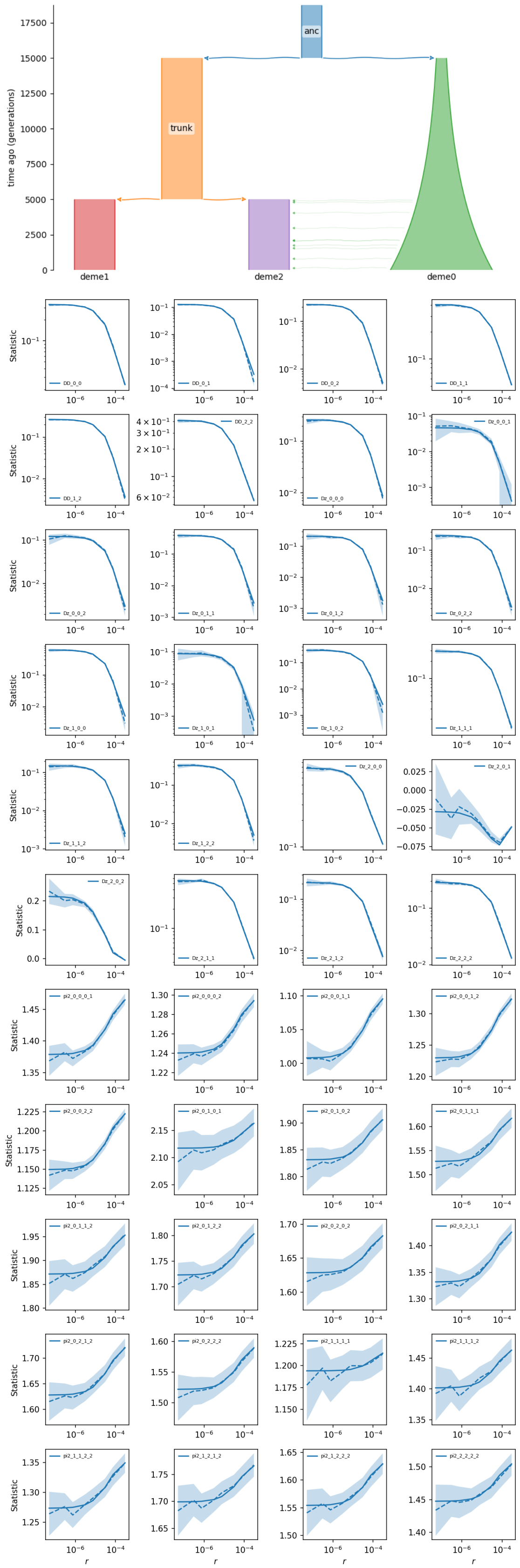


Figure S12: Demographic inference using `pg_gpu` as the `moments.LD` parser. Top: the three-population isolation model used for simulation. Bottom: LD decay curves under the inferred model (`pg_gpu`-calculated, blue; `moments`-parsed, green; truth, dashed orange) and empirical bin estimates (black points). Inference using the two parsers is indistinguishable; the drop-in [illegible]

### Extended Design and Implementation

#### Core data abstractions

Two thin wrapper classes carry data through every analysis path. The `HaplotypeMatrix` stores phased haplotypes as an $n_{\text{hap}} \times n_{\text{var}}$ 8-bit integer array (with $-1$ denoting a missing genotype), variant positions, and optional named sample sets. The companion `GenotypeMatrix` stores unphased diploid genotypes as an $n_{\text{samp}} \times n_{\text{var}}$ 8-bit unsigned-integer array—a layout chosen so that kernels whose arithmetic factorizes over the genotype tabulation $(n_{AA}, n_{Aa}, n_{aa})$ read contiguous memory. Both classes retain a single canonical buffer per object and route between CPU (`NumPy`) and GPU (`CuPy`) backings transparently via a `device` property, so the same call works whether the caller has a tree sequence (Jeffery et al. 2026) (`HaplotypeMatrix.from_ts`), a `zarr` store (`from_zarr`, supporting both `bio2zarr/VCZ` (Czech et al. 2024) and `scikit-allel` (Miles et al. 2023) layouts), or an in-memory `NumPy` array (Harris et al. 2020).

#### Streaming data abstractions

Inputs that exceed GPU memory are handled by a streaming counterpart to the in-memory matrices. The same `HaplotypeMatrix` and `GenotypeMatrix` interfaces accept a `VCZ` (Czech et al. 2024) `zarr` store and iterate over it block by block, with the on-disk variants-by-samples chunk matching the layout the fused kernels expect so each block flows through unchanged. A worker thread reads and decodes the next block while the GPU is reducing the current one, hiding I/O behind compute. Data is moved straight from disk to the GPU via `kvikio` (RAPIDS Development Team 2021) and NVIDIA's nvCOMP, with a host-side decode fallback for stores whose codec the GPU cannot decompress. Per-site, per-window, and SFS-style outputs accumulate across blocks to produce the same chromosome-wide result that the in-memory path would compute.

#### Fused kernels

Most population-genetics statistics decompose into per-site or per-pair reductions whose CPU implementations spend most of their time moving data rather than computing: each pass over the haplotype matrix reads tens of gigabytes of memory to emit a handful of summary scalars. `pg_gpu` takes advantage of this by writing *fused* kernels that read each input element once and emit *any subset* of a related family of statistics in a single pass. The entry point `windowed_analysis` accepts a list of statistic names and dispatches the request to the smallest covering set of fused kernels: a single-population kernel produces any combination of $\pi$, $\theta_W$, $\theta_H$, Tajima's $D$, Fay & Wu's $H$, the maximum DAF estimator, segregating-site counts, and singleton counts; a two-population kernel additionally produces Hudson's $F_{ST}$, Weir–Cockerham $F_{ST}$, $d_{XY}$, and $d_a$. Statistics that depend on haplotype-pattern hashing (Garud's $H_1$, $H_{12}$, $H_{123}$, $H_2/H_1$) and on scatter-add reductions (mean nSL, the DAF histogram, $\mu_{\text{SFS}}$, per-window $Z_{nS}$ and $\omega$, $\mu_{\text{LD}}$, SNP-distance moments) reuse the same window-binning infrastructure through dedicated kernel families.

##### Memory-aware chunking

A single-launch fused kernel materializes a transposed working copy of the matrix—an 8-bit integer or 32-bit float array, depending on the kernel—that can exceed GPU memory for large datasets. `pg_gpu` projects this working-set size at runtime from a free-memory query and routes the workload through a chunked variant of the same kernels whenever it would exceed 70 % of free GPU memory. The chunked path tiles the variant axis, calls the same kernels per chunk, and accumulates partial sums to the final result, producing numerically identical output to the single-pass path. This was sufficient to keep the full 10.9-million-variant Ag1000G arm on a single 80 GB A100 with no user tuning.

##### Missing data, accessibility, and span normalization

Missing data and accessibility enter the single-locus ratios through separate terms. Every diversity, divergence, and SFS estimator uses a per-site count of non-missing samples in the numerator, so allele-frequency estimates and their variance terms remain unbiased when the called-sample fraction varies across sites. For the denominator, an optional `AccessibleMask`—a boolean array over genomic coordinates, supplied directly or loaded from a BED file—replaces the nominal window width with the callable-bases count in each window. Without such a mask, gaps, low-mappability regions, and repeat-masked positions enter the denominator and silently deflate per-base values for length-normalized statistics ($\pi$, $\theta_W$, $d_{XY}$ per site) in poorly accessible regions, distorting windowed scans even when every per-site number is itself unbiased. The per-site valid-sample-count handling keeps the diversity estimator unbiased across missing-data rates from 0 to 60 % (Supplementary Figure S6). The convention matches `scikit-allel`'s windowed diversity and divergence functions, which use the same per-site valid-count term in the numerator and accept an accessibility array to mask sites that are flagged for exclusion.

##### Two-locus statistics

Two-locus (linkage-disequilibrium) statistics are the most expensive members of the catalog and the ones `pg_gpu` accelerates most, since the pairwise computation over variant sites is inherently $O(n_{\mathrm{var}}^2)$. `pg_gpu` factors every two-locus statistic through a single count-based kernel: for each pair of sites it forms the four haplotype-pair counts ($n_{AB}, n_{Ab}, n_{aB}, n_{ab}$) and the per-pair non-missing sample count, and a kernel with one thread per site pair emits any requested subset of $D^2$, $Dz$, $\pi_2$, $r^2$, $D'$, and Pearson $r$ in double precision. Windowed two-locus summaries (Kelly's $Z_{nS}$, Kim & Nielsen's $\omega$, RAiSD's $\mu_{\mathrm{LD}}$) reuse the window-binning machinery of the fused kernels; $Z_{nS}$ additionally offers a sampled-pairs estimator that trades the full $O(n_{\mathrm{var}}^2)$ sum for a sub-quadratic profile (Figure S4). The same count-based path underlies the `moments`-LD drop-in replacement (below), where the largest single speedup ($\sim$1,750×) is realized.

##### Interfacing with downstream tools

`pg_gpu` provides a drop-in replacement for the LD-parsing module of `moments` (Ragsdale and Gravel 2019): the `compute_ld_statistics` function consumes the same `VCF`/ recombination-map / population-file inputs and returns a dictionary in `moments`' canonical schema `(bins, sums, stats, pops)`, so existing infrastructure (`moments.LD.Parsing.bootstrap_data`, `optimize_log_lbfgsb`, and uncertainty quantification) accepts the output unchanged. Local PCA / `lostruct` (H. Li and Ralph 2019) is implemented as a single batched-Gram eigendecomposition over windowed views of the matrix; the subsequent inter-window MDS and corner-detection steps follow H. Li and Ralph (2019).

#### Extended feature description

The `pg_gpu` API mirrors the category organization of Table 2: each category corresponds to a Python module (`pg_gpu.diversity`, `pg_gpu.divergence`, `pg_gpu.selection`, and so on), and each statistic is a function that takes a `HaplotypeMatrix` or `GenotypeMatrix` as its first argument and returns a `NumPy` array. Population labels assigned once via `load_pop_file` are accepted by every statistic that needs them, i.e., `divergence.fst_hudson(h, "pop1", "pop2")` and `diversity.tajimas_d(h, population="pop1")` share the same calling convention. Where a statistic is defined for both phased and diploid input, the same call dispatches on container type: `selection.garud_h` computes haplotype-frequency $H$ statistics on a `HaplotypeMatrix` and the unphased diploid analogue on a `GenotypeMatrix` without separate method names. Missing data and accessibility are handled automatically.

Beyond the classic estimators, `pg_gpu` exposes the Schrider et al. (Schrider et al. 2018) distance-based two-population statistics ($Z_x$, $dd$, $dd$ rank) and the RAiSD $\mu_{\mathrm{LD}}$ component (Alachiotis and Pavlidis 2018) as ordinary functions; Patterson's $f$-statistics (Patterson, Moorjani, et al. 2012) return per-site numerators and denominators separately so that ratio-of-sums standard errors can be computed correctly with resampling. The pairwise-distance moments of Schrider et al. (2018) and the diplotype-frequency spectrum used for sweep-classifier features are also exposed directly.

The `FrequencySpectrum` class provides a direct interface to the Achaz (Achaz 2009) weighted-SFS formulation, supporting arbitrary $\theta$ estimators or neutrality tests outside the named catalog. A `FrequencySpectrum` built from a `HaplotypeMatrix` computes the SFS once on the GPU and then exposes three operations: `theta(weights)` evaluates any linear estimator $\hat{\theta} = \sum_{\xi} w_{\xi} \xi p_{\xi}$ given either a name from the built-in dictionary (`pi`, `watterson`, `theta_h`, `theta_l`, `eta1`, etc.) or a user-supplied callable that returns a weight vector; `neutrality_test` pairs any two estimators into a normalized $D$-style test statistic with a variance computed from the cached $\alpha_n, \beta_n$ coefficients of Achaz (2009), so a custom neutrality test built around an unusual weighting (e.g., higher-order powers of the frequency spectrum) is one call rather than a hand-derived variance formula; and `project` produces a hypergeometrically sub-sampled SFS at any target sample size (Gutenkunst et al. 2009), allowing comparisons across populations with different sample sizes and bridging to demographic-inference pipelines that operate on a projected SFS.

Almost every statistic in Table 2 is also accessible through `windowed_analysis`, a single entry point for genome scans that takes a list of statistic names, a window definition (in either base pairs or SNPs), and a list of populations, and returns a `pandas.DataFrame` (McKinney 2010) with one row per window. Internally, `windowed_analysis` routes the request through the fused-kernel families described above, so a typical scan that asks for $\pi$, $\theta_W$, Tajima's $D$, $F_{ST}$, $d_{XY}$, Garud's $H_{12}$, and mean nSL—seven statistics spanning four kernel families—runs in a small constant number of GPU launches over the matrix rather than seven independent passes. Local PCA / `lostruct` (H. Li and Ralph 2019) is exposed through the same interface and returns a `LocalPCAResult` with per-window eigenvalues and eigenvectors, which the companion functions `pc_dist` and `corners` transform into the inter-window distance matrix and outlier-window identifications shown in Figure 3.

General-purpose `block_jackknife` and `block_bootstrap` estimators (Busing, Meijer, and Leeden 1999; Efron and Tibshirani 1993) provide calibrated standard errors and confidence intervals for any scalar aggregate of windowed output. Both accept a tuple of (numerator, denominator) block arrays in addition to a single array, so ratio-of-sums quantities such as Patterson's $f_3$, $D$, and $F_{ST}$ are resampled with the same block assignments on the two sides of the ratio without manual bookkeeping.

Table S2: Complete catalog of statistics implemented in `pg_gpu`.

| Category | Statistic | Reference |
|---|---|---|
| Diversity | Nucleotide diversity ($\pi$) | (Nei and W.-H. Li 1979) |
| | Watterson's $\theta_W$ | (Watterson 1975) |
| | Fay & Wu's $\theta_H$ | (Fay and Wu 2000) |
| | Zeng's $\theta_L$ | (Zeng et al. 2006) |
| | Tajima's $D$ | (Tajima 1989) |
| | Fay & Wu's $H$, normalized $H^*$ | (Zeng et al. 2006) |
| | Zeng's $E$ | (Zeng et al. 2006) |
| | Zeng's $DH$ joint test | (Zeng et al. 2006) |
| | Segregating sites, singletons | (Fu 1995) |
| | Haplotype diversity, count | (Nei 1987) |
| | Expected, observed heterozygosity | (Nei 1973) |
| | Wright's $F$ | (Wright 1951) |
| | Max DAF, DAF histogram | – |
| | RAiSD $\mu_{\text{VAR}}$, $\mu_{\text{SFS}}$ | (Alachiotis and Pavlidis 2018) |
| | Diplotype frequency spectrum | – |
| Divergence | $F_{ST}$ (Hudson) | (Hudson, Slatkin, and Maddison 1992) |
| | $F_{ST}$ (Weir–Cockerham) | (Weir and Cockerham 1984) |
| | $F_{ST}$ (Nei) / $G_{ST}$ | (Nei 1973) |
| | $d_{XY}$ | (Nei 1987) |
| | $D_a$ (net divergence) | (Nei and W.-H. Li 1979) |
| | PBS | (Yi et al. 2010) |
| | $S_{nn}$ | (Hudson 2000) |
| | $G_{\min}$ | (Geneva et al. 2015) |
| | $d_d$, $d_d$ rank | (Schrider et al. 2018) |
| | $Z_x$ | (Schrider et al. 2018) |
| LD | $r$, $r^2$ | (Hill and A. Robertson 1968) |
| | $D^2$, $D_z$, $\pi_2$ | (Ragsdale and Gravel 2020) |
| | $\sigma_d^2$ (unbiased) | (Ragsdale and Gravel 2020) |
| | Kelly's $Z_{nS}$ | (Kelly 1997) |
| | Kim & Nielsen's $\omega$ | (Kim and Nielsen 2004) |
| | $\mu_{\text{LD}}$ (RAiSD) | (Alachiotis and Pavlidis 2018) |
| | Fused windowed LD | – |
| | `moments` drop-in LD parsing | – |
| Selection | iHS | (Voight et al. 2006) |
| | nSL | (Ferrer-Admetlla et al. 2014) |
| | XP-EHH | (Sabeti, Varilly, et al. 2007) |
| | XP-nSL | (Szpiech et al. 2021) |
| | $H_1$, $H_{12}$, $H_{123}$, $H_2/H_1$ | (Garud et al. 2015) |
| | EHH decay | (Sabeti, D. E. Reich, et al. 2002) |

*Continued on next page*

| **Category** | **Statistic** | **Reference** |
|---|---|---|
| SFS | Unfolded SFS, folded SFS | – |
| | Scaled SFS, folded-scaled SFS | – |
| | Joint (2-pop) SFS, folded joint SFS | – |
| | Scaled joint SFS, folded-scaled joint SFS | – |
| Admixture | Patterson's $F_2$ | (Patterson, Moorjani, et al. 2012) |
| | Patterson's $F_3$ | (Patterson, Moorjani, et al. 2012) |
| | Patterson's $D$ (ABBA-BABA) | (Patterson, Moorjani, et al. 2012) |
| | Windowed $F_3$, $D$ | (Patterson, Moorjani, et al. 2012) |
| | Block-jackknife $F_3$, $D$ standard error | (Busing, Meijer, and Leeden 1999) |
| Dim. reduction | PCA | (Patterson, Price, and D. Reich 2006) |
| | Randomized PCA | (Halko, Martinsson, and Tropp 2011) |
| | PCoA (classical MDS) | – |
| | Local PCA / `lostruct` (windowed eigendecomposition) | (H. Li and Ralph 2019) |
| | Corner detection (Welzl minimum enclosing circle) | (H. Li and Ralph 2019) |
| Resampling | Block jackknife (incl. unequal blocks) | (Busing, Meijer, and Leeden 1999) |
| | Block bootstrap | (Efron and Tibshirani 1993) |
| Generalized $\theta$ | Weighted-SFS $\theta$ estimator | (Achaz 2009) |
| | Paired neutrality tests with cached variance | (Achaz 2009) |
| | SFS projection (hypergeometric) | (Gutenkunst et al. 2009) |
| Relatedness | GRM | (Yang et al. 2011) |
| | IBS | (Purcell et al. 2007) |
| Distance dist. | Pairwise Hamming distance | – |
| | Variance of distances | (Schrider et al. 2018) |
| | Skewness of distances | (Schrider et al. 2018) |
| | Excess kurtosis of distances | (Schrider et al. 2018) |
